# Enhancer and promoter usage in the normal and failed human heart

**DOI:** 10.1101/2020.03.17.988790

**Authors:** Anthony M. Gacita, Lisa Dellefave-Castillo, Patrick G. T. Page, David Y. Barefield, J. Andrew Waserstrom, Megan J. Puckelwartz, Marcelo A. Nobrega, Elizabeth M McNally

## Abstract

The failed heart is characterized by re-expression of a fetal gene program, which contributes to adaptation and maladaptation in heart failure. To define genomewide enhancer and promoter use in heart failure, Cap Analysis of Gene Expression (CAGE-seq) was applied to healthy and failed human left ventricles to define short RNAs associated with both promoters and enhancers. Integration of CAGE-seq data with RNA sequencing identified a combined ∼17,000 promoters and ∼1,500 enhancers active in healthy and failed human left ventricles. Comparing promoter usage between healthy and failed hearts highlighted promoter shifts which altered amino-terminal protein sequences. Comparing enhancer usage between healthy and failed hearts revealed a majority of differentially utilized heart failure enhancers were intronic and primarily localized within the first intron, identifying this position as a common feature associated with tissue-specific gene expression changes in the heart. This dataset defines the dynamic genomic regulatory landscape underlying heart failure and serves as an important resource for understanding genetic contributions to cardiac dysfunction.

## INTRODUCTION

Heart failure is a leading cause of death worldwide.^1, 2^ The failed heart is characterized by reduced function, impaired filling, and altered metabolism, all of which contribute to an inability to meet the body’s demands for normal activity. Heart failure is associated with global changes in gene expression and splicing, some of which directly drive pathological and adaptive remodeling.^3^ For example, the failed heart shifts its metabolism towards glycolysis, driven in part by gene expression changes.^4, 5^ Within the sarcomere, isoform switches alter the composition of myosin heavy chain in the failed heart,^6, 7^ and alternatively spliced isoforms *TNNT2* and *TTN*, encoding troponin T and titin, respectively, directly modify contractility and compliance,^8-10^ and importantly many of these switches differ between the human and mouse heart. In addition, mutations within the coding region of many of these genes directly lead to cardiomyopathy and heart failure.^11, 12^ Although specific genetic regulatory regions have been well characterized,^13-15^ comparatively few genomewide analyses have been conducted. Genomewide epigenomic profiles have been used to infer regulatory regions in the developing mouse heart and embryonic stem cell derived-cardiomyocytes.^16, 17^ Chromatin capture integrated with CTCF binding site maps was applied to mouse cardiomyocytes subjected to pressure overload.^18^ However, much less is known about the promoter and enhancer shifts underlying human heart failure.

Gene expression is driven by transcription factor binding at promoters and enhancers, which interact in three-dimensional space to increase gene expression. Estimates of number of active heart enhancers vary from several thousand to tens of thousands depending on the approach used.^19, 20^ One assessment of active cardiac enhancers defined p300/CBP binding sites from human fetal and one adult failed heart, requiring candidate enhancer regions position > 2.5kb from any annotated transcriptional start site.^19^ This analysis suggested there were 5,000 enhancers active in fetal tissue and over 2,000 enhancers active in adult tissue, with approximately half of adult heart enhancers also active in fetal heart and underscoring the fetal re-expression program.^19^ A similar approach used normal human and mouse hearts integrating p300/CBP binding sites with H3K27ac marks and requiring candidate enhancers regions position > 1.5kb from any annotated transcriptional start site.^20^ This integrated approach described more than 80,000 potential heart enhancers.^20^ These studies provide a valuable dataset of enhancers in the normal developing and adult heart, but do not identify the genomic alterations seen in pathologic states, such as heart failure.

In addition to the use of alternative regulatory elements during heart failure, alternative promoter usage represents a distinct mechanism to regulate gene expression. Typically driven by the inclusion of alternative first exons, alternative promoters are estimated to affect 30-50% of human genes.^21, 22^ Alternative promoters may affect the amino-terminal amino acids of proteins and/or the 5’UTR of transcripts, both of which can mediate functional consequences. Alternative promoters can also influence the effect of genetic variants on protein function and thus are vital for accurate variant effect predictions.^23^ Despite the potential of broad proteome differences due to alternative promoter usage, a genomewide view of promoter and enhancer shifts in human heart failure is lacking.

Next generation sequencing technologies including Cap-Analysis of Gene Expression (CAGE) have made it possible to assay transcriptome usage by determining RNA transcriptional start sites at single base pair resolution.^24^ Enhancer regions are transcribed into low-abundance enhancer RNAs (eRNAs) in a bidirectional pattern,^25, 26^ contrasting with the unidirectional transcriptional expression seen near gene promoters, which produce high-abundance signals. Because of the precision with which these RNAs can be mapped, it is possible to accurately map enhancer and promoter signals at high resolution and without using blanket arbitrary filters that ultimately limit identification. For example, enhancers throughout the genome can be identified, as this analysis does not require removal of promoter-proximal intervals.

To define alternative promoter and enhancer use in heart failure, we generated CAGE sequence datasets from healthy and failed human left ventricles. CAGE sequencing information was integrated with RNA sequencing from these same samples, to improve sensitivity in detecting low-abundance eRNAs and rarely used gene promoters. We relied on a no-amplification non-tagging CAGE sequencing protocol, which allows for more robust and less biased detection of transcriptional start sites.^24^ These data identified unique signatures of housekeeping and tissue specific promoters, as well as a pattern of enhancers within first introns that regulate tissue specific expression. In addition to identifying differential enhancer use in heart failure, we cataloged 129 genes with differential promoter usage in heart failure. These alternative promoters have the potential to encode proteins with unique amino-termini, highlighting potential protein composition shifts in the failed heart.

## RESULTS

### CAGE sequence clusters identify promoters and enhancers of the left ventricle

CAGE sequencing identifies promoters and enhancer regions. Given the known gene expression differences that characterize failed hearts, we generated CAGE datasets from left ventricle (LV) taken from three healthy and four failed hearts. Healthy LV samples were those acquired but not used for transplant due to age or other incompatibility. Failed hearts were obtained at the time of transplant from patients with a range of primary mutations and ages (**Table 1**). Each library was sequenced to high depth and libraries demonstrated comparable alignment rates (**Supplemental Table 1**). To generate a comprehensive list of all potential promoters and enhances, the initial analysis included the combination of healthy and failed hearts. CAGE sequence analysis identified 23,676 unidirectional sequence clusters, indicative of promoter regions, and 5,647 bidirectional sequence clusters, indicative of enhancer regions.

**TABLE 1.** Left Ventricle Tissue Source Demographics, Phenotypes, and Mutations.

| **Sample** | **Primary phenotype** | **Additional phenotypes** | **Sex** | **Race** | **Age** | **Primary gene mutation(s)** |
| --- | --- | --- | --- | --- | --- | --- |
| Healthy 1 | Healthy | - | M | Caucasian | 62 | N/A |
| Healthy 2 | Healthy | - | M | Caucasian | 47 | N/A |
| Healthy 3 | Healthy | - | F | Caucasian | 76 | N/A |
| Heart Failure 1 | Cardiomyopathy | Ventricular Tachycardia | M | Caucasian | 20 | *TPM1* D230N |
| Heart Failure 2 | Cardiomyopathy | - | M | Hispanic | 16 | *TTN* c.42521-5 C>G,<br>*TNNT2* K210del |
| Heart Failure 3 | Cardiomyopathy | Becker Muscular Dystrophy | M | Caucasian | 54 | *DMD* IVS +1 G>T |
| Heart Failure 4 | Cardiomyopathy | Limb Girdle Muscular Dystrophy | F | Caucasian | 26 | *LMNA* c.1142-1157+1del17 |

Unidirectional CAGE sequence clusters were annotated using Ensembl designations to map their position relative to annotated genes. Unidirectional sequence clusters that mapped ±100bp of transcriptional start sites constituted 70.1% of the clusters, consistent with their putative role as promoters (**Figure 1A**). An additional 8.1% of unidirectional clusters, mapped between 100 and 1000bp upstream of transcriptional start sites. The remaining 21.8% of sequence clusters mapped to 5’UTR, 3’UTR, exons, introns or intergenic regions.

**Figure 1.**
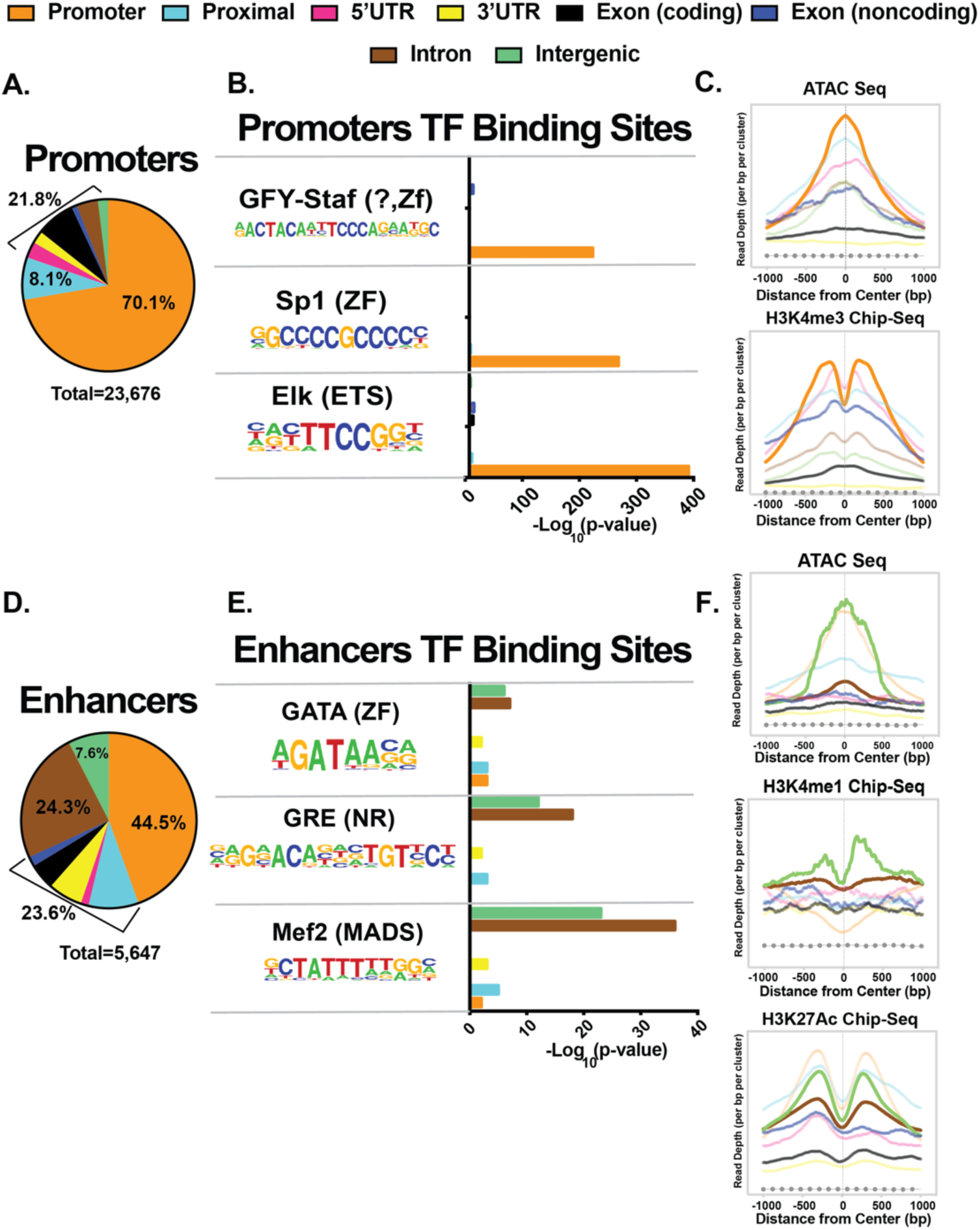
CAGE sequencing identified promoters and enhancers active in human left ventricular tissue. **A.** Shown is the distribution annotation classes of unidirectional CAGE clusters. **B.** Enrichment values for three promoter-bound transcription factors in unidirectional clusters from all annotation classes. **C.** Left ventricle signals of open chromatin (ATAC-seq) and a promoter-associated histone mark (H3K4me3) across unidirectional clusters from all annotation classes. **D.** Venn diagram indicating the distribution annotation classes of bidirectional CAGE clusters. **E.** Enrichment values for three cardiac transcription factors in bidirectional clusters from all annotation classes. **F.** Left ventricle signals of open chromatin (ATAC-seq) and two enhancer-associated histone marks (H3K4me1 and H3K27Ac) across bidirectional clusters from all annotation classes. Dashed lines in **C** and **F** represent signals from genomic regions created by scrambling the location of unidirectional and bidirectional clusters, respectively. *TF*, transcription factor.

We next analyzed clusters for their transcription factor binding motif composition. The 70.1% of clusters mapping within 100bp of transcriptional start sites were highly enriched for GFY-Staf, Sp1, and Elk/ETS binding motifs, which are known promoter binding transcription factors.^27^ Clusters mapping into other gene regions showed minimal enrichment of these motifs (**Figure 1B**). To provide additional support for the promoter-enriched sequence clusters, ATAC sequencing and H3K4me3 ChIP-seq datasets were compared; **Supplemental Table 2** provides source information on comparison datasets. Clusters overlapping promoters overlapped considerably with ATAC-seq and H3K4me3 ChIP-seq signals, indicative of open chromatin and active promoter regions (**Figure 1C**). These clusters also showed high CTCF and Pol2A binding as well as a reduction of H3K4me1 histone modifications, consistent with their role as promoters and not enhancers (**Supplemental Figure 3**). The bimodal shape of histone methylation patterns is consistent with open chromatin signals being flanked by promoter histone marks. Taken together, these data identify unidirectional CAGE sequence clusters as bearing the genomic signatures of active promoters.

The bidirectional CAGE clusters were similarly annotated with Ensembl designations. Only 44.5% of bidirectional clusters mapped ±100bp within transcriptional start sites. In contrast to unidirectional clusters, 24.3% of clusters mapped to gene introns and 7.6% were intergenic (**Figure 1D**). When analyzed for known transcription factor motifs, intergenic and intronic clusters showed enrichment of GATA, GRE, and MEF2 motifs, essential transcription factors for cardiomyocyte specification and maintenance (**Figure 1E**).^28, 29^ Intergenic bidirectional clusters showed enrichment of open chromatin signals (ATAC-seq), H3K4me1, and H3K27Ac histone modifications in human left ventricles. Intronic clusters also showed a similar pattern, but with lower magnitude (**Figure 1F**). The intergenic and intronic bidirectional clusters showed enrichment of CTCF and Pol2A binding as well as reduced H3K4me3 modifications (**Supplemental Figure 3**). These patterns are highly consistent with the role of bidirectional CAGE sequence cluster regions as being enhancers, rather than promoters.

### CAGE sequence-defined promoters show shape divergence

Mammalian promoters initiate transcription across broad or narrow genomic regions and correlate with distinct transcriptional regulatory mechanisms.^30^ We evaluated cardiac promoters for these two major types of promoters by calculating the interquantile range (IQR) of promoter CAGE clusters by determining the base pair distance between 10% and 90% of a promoter’s total signal. We observed the expected two distinct populations, defined as sharp (IQR < 10bp) and broad promoters (IQR ≥ 10bp) (**Figure 2A**). Subjecting these two types of promoters to gene ontology analysis, we observed that broad promoter genes were those associated many cellular functions, while genes regulated by sharp (narrow) promoters were significantly enriched for muscle and cardiac gene ontology terms (**Figure 2B**). Thus, tissue specific genes important for left ventricular specification and function were more likely to have sharp promoters.

**Figure 2.**
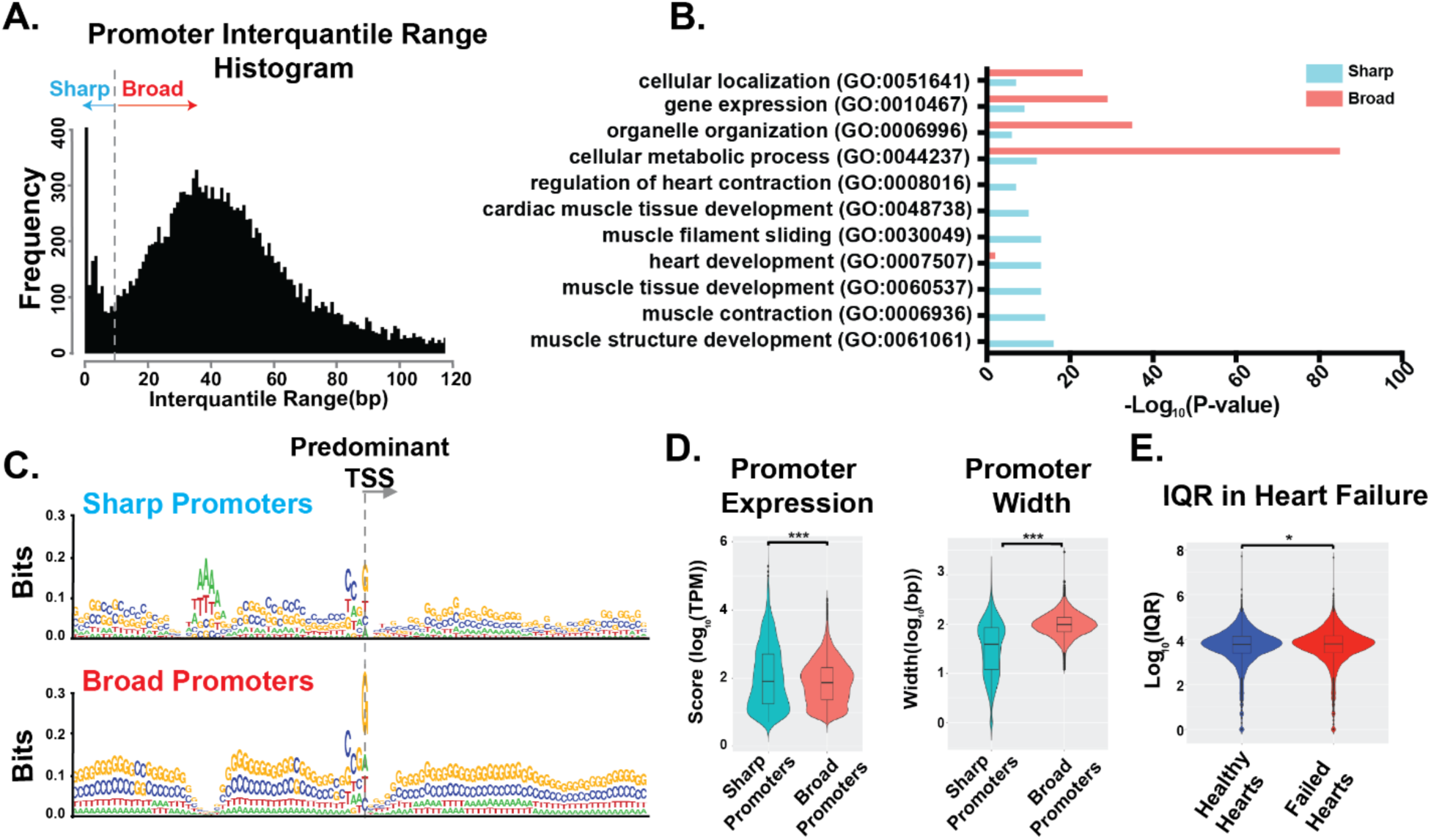
Sharp and broad transcriptional start sites in cardiac promoters. **A.** Histogram of interquantile range (number of basepairs between 10% and 90% of a total signal from a given promoter) of all promoters revealed sharp and broad promoter classes. **B.** Gene ontology analysis of genes driven by sharp or broad promoters indicates broad promoters have housekeeping functions while sharp promoters are found across all gene ontology categories including tissue specific genes. **C.** Relative nucleotide compositions of the upstream and downstream sequences from the transcriptional start site for sharp and broad promoters. **D.** Violin plots comparing the expression level and width of sharp and broad promoters. **E.** Violin plot comparing the interquantile range of promoters in healthy and failed hearts. Significance determined by two-tailed nonparametric Mann Whitney Test (p ≤ 0.05(*), ≤ 0.005(**), ≤ 0.0005 (***)). *TSS*, transcriptional start site. *IQR*, interquantile range. *bp*, basepair.

Sharp and broad promoters also demonstrated differential enrichment of upstream sequence DNA-binding motifs. Sharp promoters had TATA motifs at positions 30-33 upstream of the predominant transcriptional start site, representing canonical TATA boxes. Broad promoters were devoid of TATA motifs, but did show enrichment of GC nucleotides, which likely represents CpG-islands.^31^ Both classes of promoters showed a preference of transcription initiation at a G nucleotide and an upstream CC sequence at the −2 and −3 positions (**Figure 2C**). Sharp, tissue specific promoters were also more highly expressed compared to broad promoters, and this observation was driven by a smaller population of very highly expressed sharp promoters (**Figure 2D**). We compared promoter shape between healthy and failed hearts and found a modest but significant genomewide increase in promoter IQR in failed hearts (**Figure 2E**).

### Intronic Enhancers map within the first Intron

A large proportion of bidirectional CAGE clusters are located within gene introns. Intronic clusters shared transcription factors and epigenetic marks with intergenic CAGE clusters, suggestive of their roles as enhancers (**Figure 1E**). While a typical human gene contains on average 7.8 introns,^32^ we observed that the majority (69%) of intronic enhancers in this dataset mapped to the first intron (**Figure 3A**). First intron enhancers generated more eRNA than enhancers in other introns, but were not wider and did not differ in their balance of bidirectionality (**Figure 3B**). Notably, all intronic enhancers mapped within genes enriched for cardiac and muscle gene ontology terms (**Figure 3C**), suggesting the importance of first intron enhancers for determining tissue specificity.

**Figure 3.**
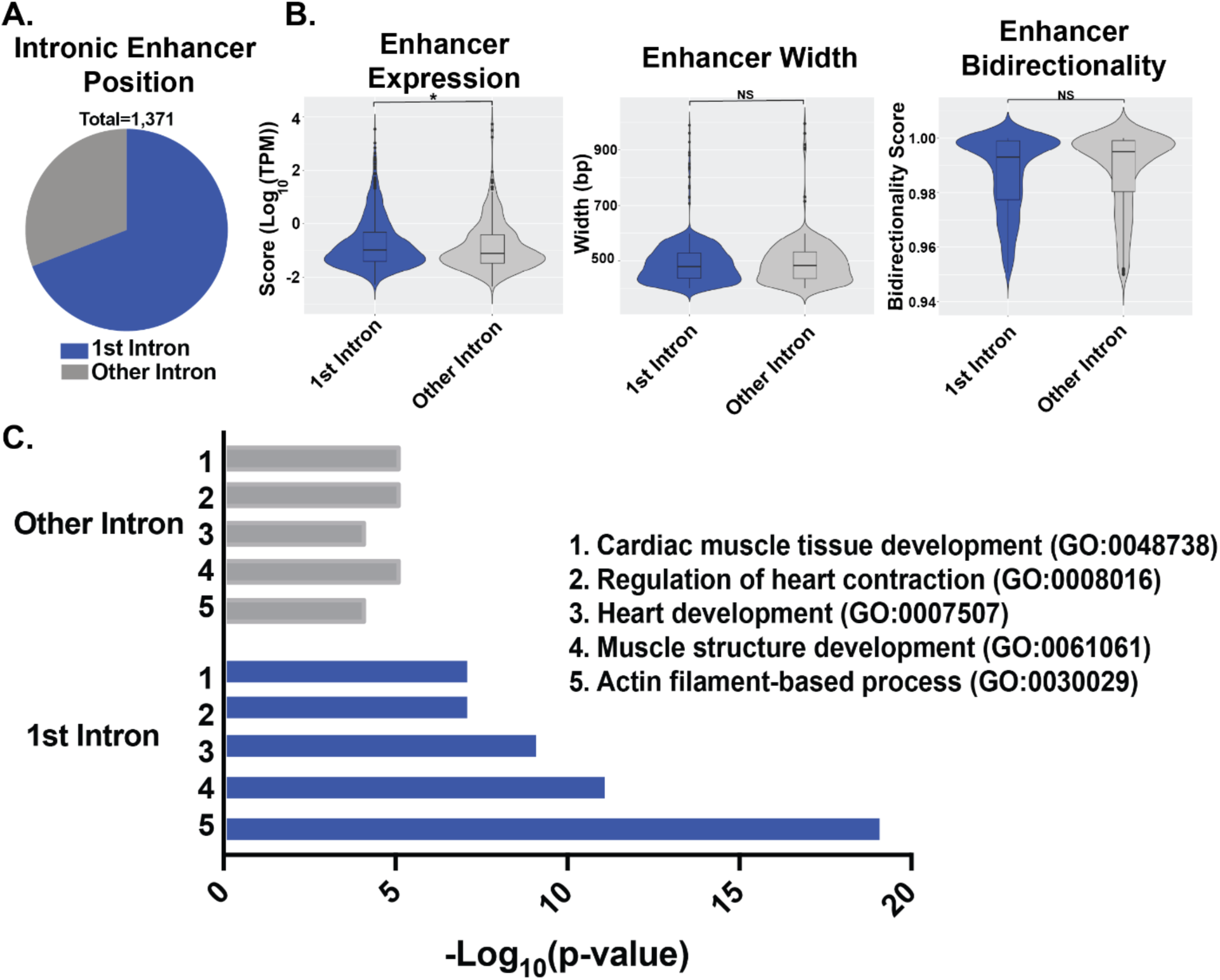
The majority of intronic cardiac enhancers localize to the first intron. **A.** Venn diagram demonstrating that ∼70% of intronic enhancers were within the first intron of an overlapping transcript. **B.** Violin plots comparing enhancer expression levels, enhancer width, and enhancer bidirectionality score between enhancers in the first intron and enhancers in other introns. **C.** Gene ontology analysis of genes with first intron enhancers and genes with other intron enhancers indicates tissue specific genes are more likely to have first intron enhancers. Significance determined by two-tailed nonparametric Mann Whitney Test (p ≤ 0.05(*), ≤ 0.005(**), ≤ 0.0005 (***)).

### Correlation of CAGE-sequencing and RNA-sequencing

RNA sequencing was carried out on the same left ventricles and compared to CAGE sequencing. Since CAGE sequencing quantifies promoter expression, it also can measure overall gene expression, and this was consistent with the tight correlation between CAGE sequencing and RNA sequencing (**Figure 4A**). Additionally, we assessed correlations between pairs of samples. In general, healthy hearts correlated best with other healthy hearts, and failed hearts compared best with failed hearts. The RNA sequence and CAGE sequence expression estimates were most correlated for matched samples except for Failed Heart 4, which likely reflects the lower CAGE sequence read depth in this sample (**Figure 4B**). Gene expression differences were identified using both CAGE and RNA sequence datasets. RNA-seq identified more upregulated and downregulated genes, and approximately half of the genes identified by CAGE-seq were also identified by RNA-seq (**Figure 4C**). Gene ontology analyses on differentially expressed genes were similar in both sequence datasets. Genes associated with developmental pathways and extracellular matrix organization were upregulated in heart failure while genes associated with catabolism were downregulated in heart failure (**Figure 4D**), consistent with prior reports.^3, 4^

**Figure 4.**
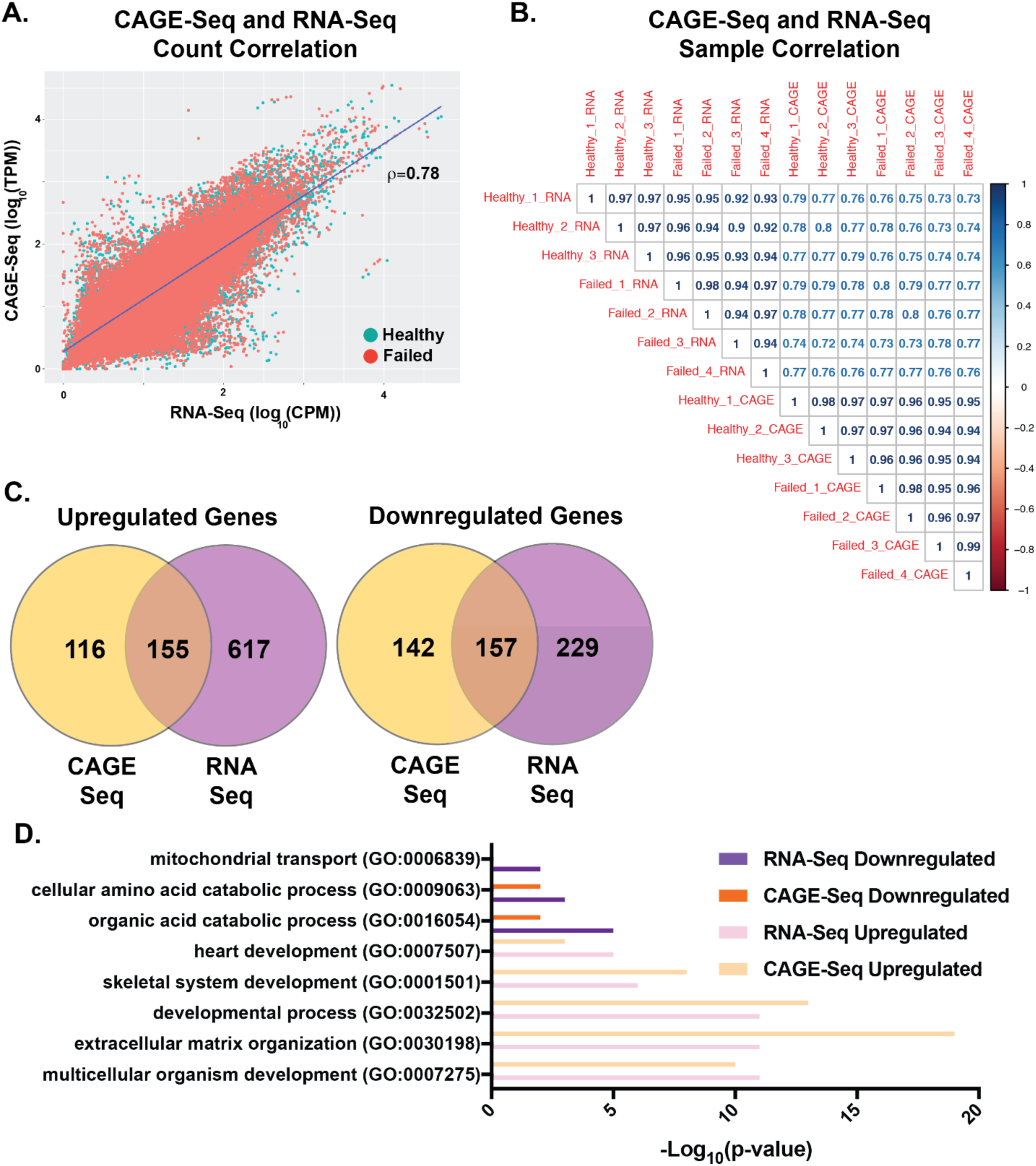
Comparison of CAGE-seq and RNA-seq gene expression levels. **A.** Scatter plot of CAGE-seq gene expression values versus RNA-seq expression values for healthy (teal) and failed (red) samples demonstrating tight correlation. **B.** Sample level correlation matrix of Spearman’s correlation coefficient of genome-wide gene expression levels. **C**. Venn diagrams displaying the number of differentially upregulated and downregulated genes determined by CAGE-seq and RNA-seq. **D.** Gene ontology analysis of genes identified as differentially upregulated or downregulated by CAGE-seq and RNA seq.

### CAGE sequencing-defined enhancer regions validated by other enhancer datasets

Of the ∼1,800 enhancer regions identified by CAGE sequencing, data was available from 45 of these in the Vista Enhancer Browser, a database of enhancers tested in an *in vivo* reporter assay using transgenic mouse embryos.^33^ Of the 45 present in Vista, 31 (70%) demonstrated reproducible activity in the developing mouse heart) (**Supplementary Figure 4A**). We also compared CAGE sequence-predicted enhancers to those predicted by H3K27Ac and p300 ChIP sequencing from developing and adult human and mouse tissues.^20^ CAGE-sequence defined enhancers showed significantly higher overlap to H3K27Ac/p300 ChIP regions compared to length-matched scrambled control regions (**Supplementary Figure 4B**). An additional study reporting H3K27Ac ChiP-seq data from healthy and failed human hearts was similarly compared and showed significant overlap (**Supplementary Figure 4C**).^34^ Finally, we compared CAGE sequence-defined enhancer predictions from the FANTOM consortium, which used CAGE sequencing across many tissues to define enhancers.^35^ The CAGE sequencing in this current study showed significant overlap with FANTOM predictions with the highest percentage of overlap for ubiquitous enhancers but also overlap with left ventricle predicted enhancers (**Supplementary Figure 4D**). Many enhancers described here were not present in FANTOM predictions because of higher depth sequencing in our study (**Supplementary Table 1**). Taken together, the intersection of these orthogonal datasets with CAGE sequence data corroborate the notion that we have uncovered cardiac enhancers both in healthy hearts and in heart failure.

### Alternative promoter usage in heart failure

In the LV, 3,032 (23%) expressed genes have evidence for more than one promoter (**Figure 5A**). We used CAGE sequencing data to calculate the percentage of total transcripts coming from each promoter in multi-promoter genes. We compared the average percent-usage of each promoter in healthy and failed hearts and found 609 promoters in 325 genes with a shift ≥10% (**Figure 5B**). Of these, 149 promoters in 124 genes occurred after the exon containing the start codon, indicating the potential to alter the amino-terminal amino acid sequence of the resulting protein (**Figure 5C**). Of the 124 genes identified as having alternative promoters that occur after start codons in heart failure, many are associated with sarcomere regulation or muscle structure development, including *TNNT, MYOT*, and *SPEG.* We annotated a significant promoter switch in *PRKAG2*, a gene linked to hypertrophic cardiomyopathy and critical to heart metabolism.^36-38^ Three major *PRKAG2* promoters were identified, encoding three different isoforms-γ2a, γ2-3b, and γ2b **(Figure 5D)**. In healthy hearts, the relative expression of these three transcripts is 53% γ2b, 28% γ2-3b, and 17% γ2a. In heart failure, these percentages significantly shift with 29% γ2b, 59% γ2-3b, and 10% γ2a isoform (**Figure 5E**). Notably, the γ2-3b isoform encodes a unique 32 amino acid sequence at the amino-terminus (**Figure 5F**).

**Figure 5.**
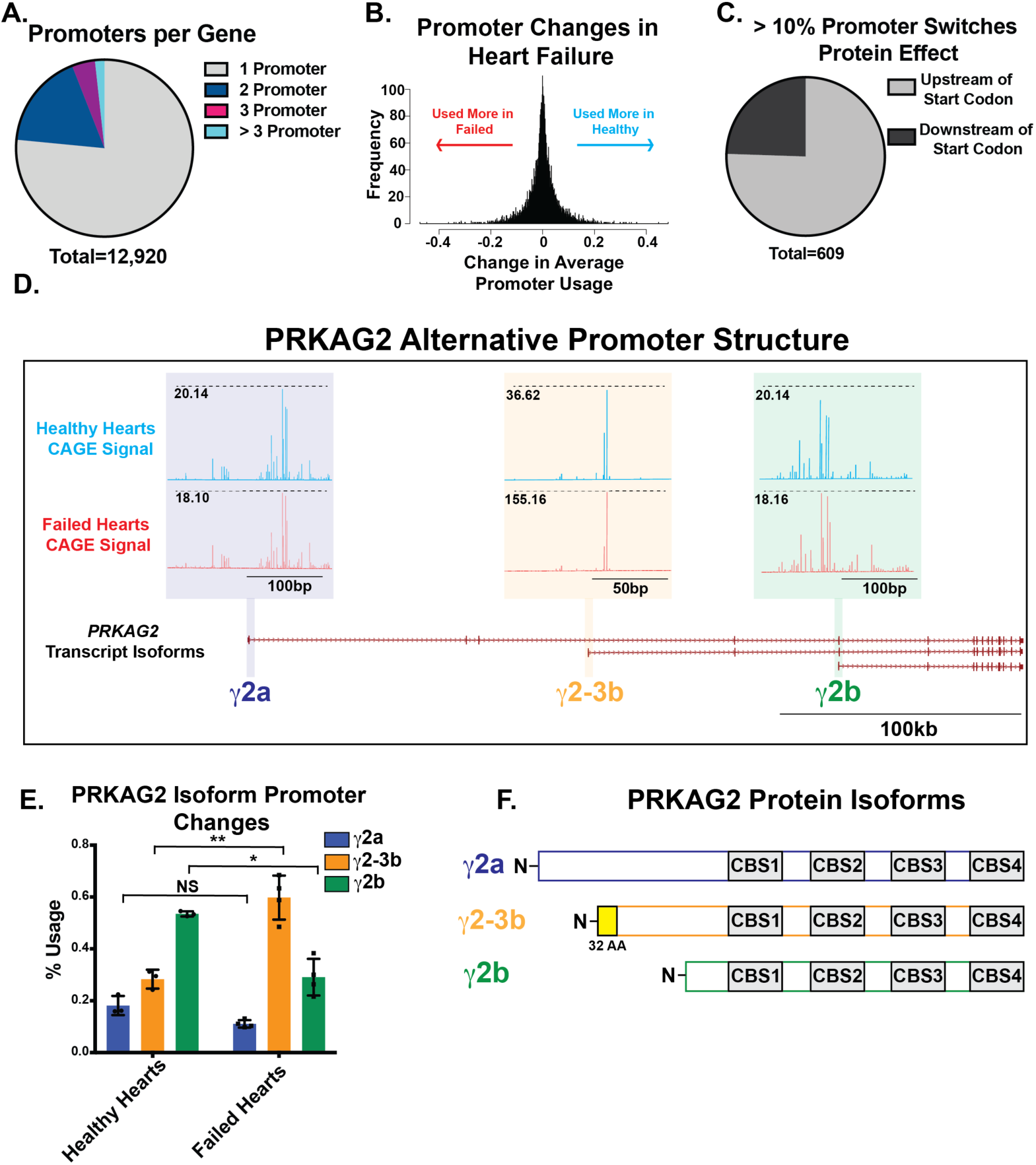
Heart failure was associated with significant changes in left ventricle promoter usage. **A.** Venn diagram of the number of left ventricle promoters per gene determined by CAGE-seq. **B.** Histogram displaying the distribution of average promoter percent usage changes in heart failure. The x-axis represents the difference between a promoter’s average percent usage in three healthy left ventricles and four failed left ventricles. Left-shifted promoters make up a greater percentage of gene expression in failed ventricles and right-shifted promoters are a greater percentage in healthy ventricles. **C.** Venn diagram of the relationship to an overlapping transcript’s start codon for promoters that undergo a ≥ 10% shift in heart failure. **D**. Genome browser representation of the alternative promoter structure of the *PRKAG2* gene. Three known isoforms of *PRKAG2* are represented at the bottom. Above the promoter of each transcript, the CAGE-seq signal for healthy (blue) and failed (red) hearts is shown and the scales of each representation are indicated in black. **E.** Quantification of the CAGE-seq signals shown in D indicating the promoter percent usage of each isoform in healthy and failed hearts. Significance determined by a two-tailed Student’s t-test. **F.** Schematic of the predicted amino acid sequences translated from each *PRKAG2* isoform. (p ≤ 0.05(*), ≤ 0.005(**), ≤ 0.0005 (***)). *PRKAG2*, Protein Kinase AMP-Activated Non-Catalytic Subunit Gamma.

### Enhancer usage shifts in heart failure

The CAGE sequence analysis identified ∼1800 enhancer regions actively transcribed in human left ventricle (**Figure 1A**). Multidimensional scaling of normalized expression levels showed an overall similar profile of enhancer usage in healthy heart samples, but more disparate enhancer usage across the four failed hearts (**Figure 6A**). Comparing enhancer usage across heathy and failed hearts reveals 264 enhancers changing significantly in heart failure (raw p-value ≤ 0.05). To assess whether differential enhancer transcription was associated with differential transcription factor binding site profiles, we compared transcription factor motif instances across enhancers in healthy and failed hearts. We found that motifs for SMAD2, NFIX, NFAT, TCF7L2, ZNF740, and AR were enriched in enhancers that change in heart failure. SMAD2, NFIX, TCF7L2, and AR motifs were found more in downregulated enhancers. While NFAT and ZNF740 motifs were found more in upregulated enhancers. RNA-sequencing demonstrated that *SMAD2* and *NFAT5* were significantly upregulated in heart failure (**Figure 6C**). **Figure 6D** illustrates alternative enhancer use within the first intron of *TRPM7*, which encodes the transient receptor potential cation channel subfamily M member, a gene implicated in ischemic cardiomyopathy and cardiac rhythm.^39, 40^ This intronic enhancer showed significantly lower eRNA expression in heart failure (**Figure 6E**), concomitant with a significantly lower expression of *TRPM7* in failed hearts (**Figure 6F**).

**Figure 6.**
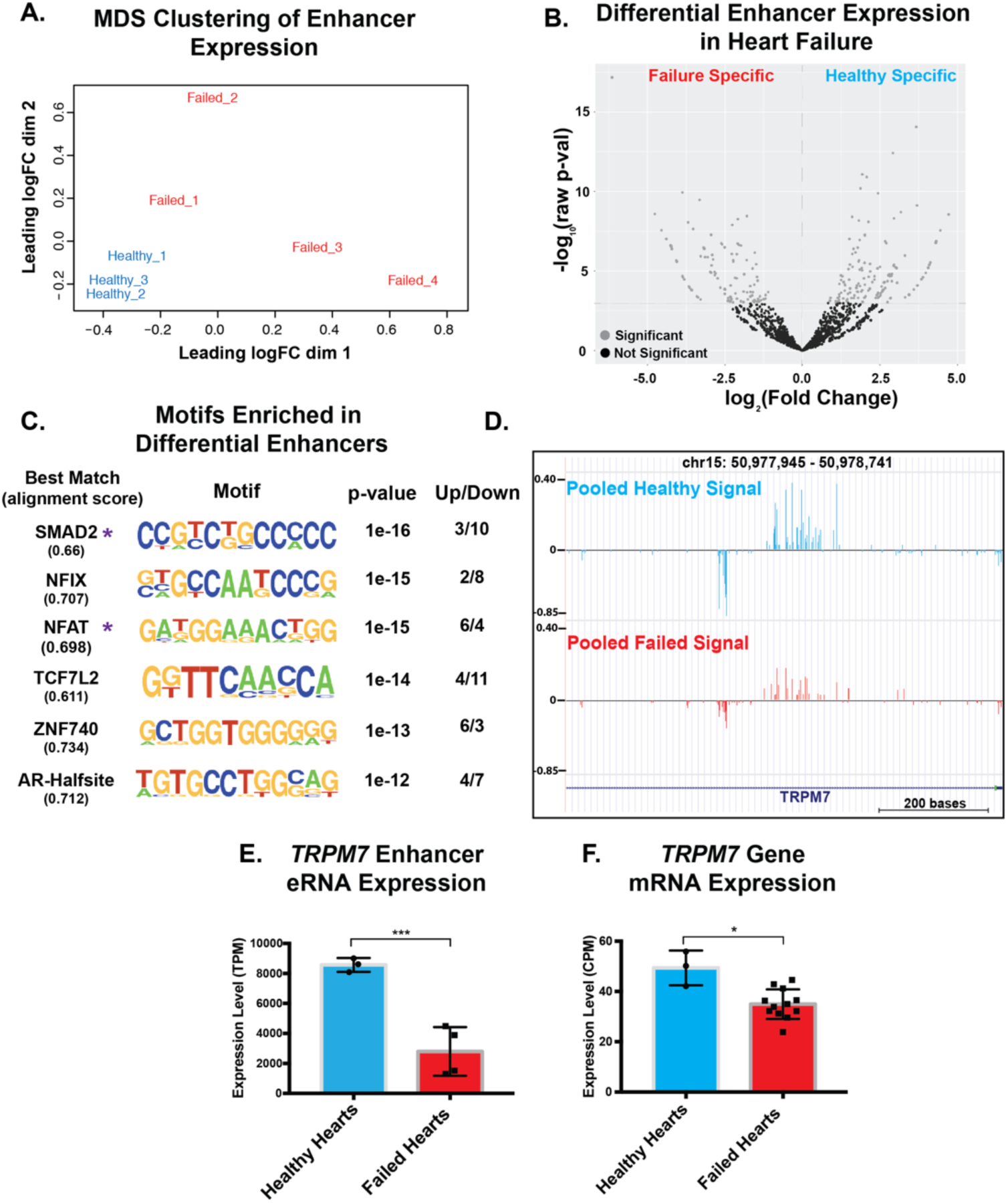
CAGE-Seq detected differential left ventricle enhancer usage in heart failure. **A.** Multidimensional scaling plot of enhancer expression levels in healthy and failed left ventricle samples. **B.** Volcano plot indicating the results of differential enhancer usage analysis. Differential enhancers are in light gray. Left shifted enhancers are expressed higher in failed hearts and right shifted enhancers are expressed higher in healthy hearts. **C.** *De novo* transcription factor motif enrichment analysis comparing differentially changed enhancers to unchanged enhancers. The best match of enriched motifs is listed to the left. A purple asterisk indicates that the matching transcription factor was differentially expressed by RNA-seq. Up/down indicates the instances of the identified motif in upregulated and downregulated enhancers, respectively. **D**. Genome browser representation of a differentially expressed enhancer within the first intron of the *TRPM7* gene. The gene annotation is at the bottom and the healthy and failed CAGE-seq signals are graphed above on the same scale. **E**. Quantification of the healthy and failed CAGE-seq signals for the intronic *TRPM7* enhancer in D. **F**. Quantification of *TRPM7* overall gene expression by RNA-seq. Significance determined by EdgeR using a generalized linear model approach. (p ≤ 0.05(*), ≤ 0.005(**), ≤ 0.0005 (***)). *FC*, fold change. *TRPM7*, transient receptor potential cation channel subfamily M member 7.

## DISCUSSION

### Defining the promoterome of the human LV in health and disease

Using CAGE-sequencing, we provide the most detailed genome-wide analysis of promoter usage in human left ventricle to date. The majority of CAGE sequence clusters overlapped the promoter regions of known annotated RNAs. Additionally, clusters mapping to promoter regions were enriched for promoter-bound transcription factors and epigenetic modifications compatible with their annotation as promoters. In total, we report ∼17,000 high likelihood promoters active in the adult human heart. We observed two major promoter types, the sharp TATA-box-associated and the broad CpG island-associated.^31^ Sharp promoters had single or a few transcriptional start sites and were linked to highly-expressed, tissue-specific genes like *MYH7, TTN*, and *MYL2*. Broad promoters had a wider distribution of transcriptional start sites and included both housekeeping and some tissue specific genes. We observed an increase in genome-wide promoter width in failed left ventricles, suggesting a loss of tight regulation of transcriptional start sites. This widening of promoters may reflect epigenetic modifications or transcriptional factor profile differences, both of which are known to occur in heart failure and hypertrophy.^41, 42^

### Alternative promoter usage in heart failure

The CAGE sequence data indicate that ∼20% of genes active in the human left ventricle have multiple active promoter regions, correlating well with previous estimates from different cell types.^43, 44^ Promoter switches can generate unique 5’UTRs without changing the resulting protein, or result in novel amino-termini if the switch activates an alternative promoter downstream of the translation start site. Altering the 5’UTR may affect translational efficiency, imparting developmental and tissue specificity.^45^ We demonstrated a significant shift in promoter usage in the *PRKAG2* gene. Mutations in *PRKAG2* have been linked to hypertrophic cardiomyopathy and arrhythmias.^36^ *PRKAG2* is highly expressed in the heart, and it was reported that the γ2b-3b isoform was most highly expressed in the heart.^46^ In healthy left ventricles, we found that ∼55% of transcripts originated from the γ2b promoter and ∼35% originate from the γ2-3b promoter. In failed left ventricles, ∼60% of *PRKAG2* transcripts represent the γ2b-3b isoform, which has a unique 32 residue amino-terminal domain that may affect the cellular localization of *PRKAG2*. Data from a yeast two-hybrid screen indicated that the amino-terminus of the longer γ2b, which shares some homology with the γ2b-3b isoform, interacts with troponin I.^47^ Upregulation of the γ2b-3b isoform in failed hearts may also confer the cardiac-specific phenotype of *PRKAG2* mutations. This data indicates that alternative promoter usage may have broad implications on the cellular proteome.

### Enhancer enrichment in the first introns

A significant portion of CAGE-defined enhancers were located within intronic regions of overlapping annotated transcripts, showing similar epigenetic modifications and transcription factor motifs to intergenic enhancer regions. The majority of intronic enhancers were located in the first intron. Genes with highly expressed first intron enhancers included *MYLK3, TPM1, SLC8A1*, and *FLNC*, which are all genes important for cardiac performance. Classic intronic enhancers are have been described in mammalian systems.^48, 49^ A study of copy number variants within intronic regions saw significant correlations between intronic deletions and gene expression in lymphoblastoid cell lines.^50^ Additionally, a study using histone modifications to identify enhancer regions active in stem-cell derived cardiomyocytes identified a large proportion of enhancers within the first intron.^16^

### Differential enhancer usage in heart failure

Heart failure is associated with both structural and transcriptional changes.^3, 51^ We found that enhancer usage was more variable in failed ventricles, which may indicate genome-wide dysregulation of gene expression. *SMAD2* binding motifs were enriched in differentially used heart failure enhancers, and this is highly consistent with the known upregulation of TGF-β signaling in failing hearts.^52, 53^ The enrichment of this motif in differential enhancers may be due to increased TGF-β/SMAD activation in cardiomyocytes and/or a larger proportion of cardiac fibroblasts in the failed left ventricle tissues. *SMAD* motifs were found more often in downregulated enhancers, implying a repressive role for TGF-β/SMAD signaling in heart failure. We highlighted a specific differential enhancer located within the first intron of the *TRPM7* gene. *TRPM7* encodes kinase domain-containing cation channel. Deletion of *Trpm7* in mice disrupts cardiac automaticity and causes cardiac hypertrophy and fibrosis.^40, 54^ In ischemic cardiomyopathy, *TRPM7* was significantly down regulated in the left atria and ventricle.^39^ However, in non-ischemic dilated cardiomyopathy, *TRPM7* was shown to be upregulated in patients with ventricular tachycardia.^55^ We found a larger magnitude reduction in enhancer eRNA levels compared to reduction in mRNA levels, and this may indicate that additional *TRPM7* enhancer regions are active. Our findings support a reduction in *TRPM7* in the setting of end stage heart failure.

### Study Limitations and Conclusions

This study used CAGE sequencing to define a broad spectrum of cardiac promoters and enhancers, with focus on their differential use in heart failure. We observed variability in differential promoter and enhancer usage in failed heart, as the normal control hearts showed tighter correlations. This variability may reflect the end stage process of heart failure. While a larger dataset may be more revealing, the diversity of response in the failed hearts mirrors what has been observed when RNA sequencing was used to define transcripts produced for *TTN*, a large gene that has been examined in multiple failed hearts.^56, 57^ The wide array of transcripts produced from even this single gene may underscore that a lack of uniform response itself could contribute to heart failure.

## METHODS

### Materials, Code and Data Availability

All scripts used in this analysis are uploaded to the github page (in progress). Sequence data has been uploaded to the NCBI-GEO under accession number (in progress).

### RNA-Extraction, Library Preparation, and Sequencing

Healthy and failed left ventricle samples were obtained from failed transplants or as discarded tissue, respectively. Living subjects provided consent. Healthy left ventricular samples were obtained from hearts provided by the Gift of Hope of Illinois and were found to be unsuitable for transplant due to age or prior cardiac surgeries. All patients were declared to be brain dead as the result of cerebral hemorrhage and familial consent was obtained for organ use in research. Tissues were snap-frozen in liquid nitrogen and stored at −80°C until use. Approximately 50mg of frozen tissue was ground into a fine powder using a mortar and pestle under liquid nitrogen. Ground powder was added to 1mL TRIzol (Invitrogen) containing 250ul of silica zirconium beads. Samples were placed in a bead homogenizer for 1 minute, allowed to cool on wet ice, and centrifuged at 12,000xg to remove any unhomogenized tissue pieces followed by chloroform extraction. Phases were separated by centrifugation and the upper aqueous layer was added to fresh 70% ethanol. The RNA-ethanol mix was used as in input to the Aurum Total RNA Mini Kit (BIORAD). RNA was isolated (including on-column DNAse digestion) following manufacturer’s instructions. Concentration was measured using a NanoDrop spectrophotometer and quality was assessed using an Aligent Bio analyzer. Only RNA extractions with RIN values ≥7 were used. If necessary, RNA extraction was repeated until ∼10μg of RNA was obtained.

Custom nAnT-iCAGE-seq (no-Amplification-no-Tagging Illumina Cap Analysis of Gene Expression libraries) libraries were prepared by DNAFORM (Japan) following a previously-described protocol ^24^. Briefly, 5μg of RNA was reverse transcribed using random primers. 5’-methyl-caps were biotinylated and enriched using streptavidin beads. cDNA was released and sequencing adapters were added using blunt-ended ligation. Following second strand synthesis, the libraries were quantified using qPCR. ∼50pM of pooled libraries were loaded into an entire run on the NextSeq 500 (Illumnia) to yield ∼400 million total 75bp single end reads (**Supplemental Table 1**).

RNA-seq libraries were prepared using the TruSeq mRNA-seq library preparation kit (Illumina). Libraries were pooled in equimolar amounts and loaded on the HiSeq 4000 (Illumina) to generate ∼40 million 150bp paired-end reads/sample.

### CAGE-Seq Alignment and Clustering

Raw CAGE-seq reads were checked with fastQC(v0.11.5) and aligned to the human genome (hg19) using STAR (v2.5.2) with default settings.^58^ Uniquely aligning reads were inputted into CAGEr and converted into quantified CAGE transcriptional start site (CTSS) coordinates with removal of first G nucleotide mismatches.^59^ CTSS coordinates and counts were outputted as bigwig files for input to CAGEfightR.^60^ CTSSs from mitochondrial chromosomes and CTSSs only present in a single sample were removed. We clustered CTSSs from all samples into unidirectional and bidirectional clusters. CTSS’s were required to have ≥5 pooled counts be included in a unidirectional cluster and all CTSSs within 20bp were merged into the same cluster. Unidirectional clusters also were required to have >1 TPM (tag per million) in at least 2 samples. Bidirectional clusters were required to have a balance score ≥0.95 and a 200bp window on either side of the midpoint was used to quantify each cluster, as described in.^35^ Bidirectional clusters were also required to be bidirectional in at least one sample and have ≥ 2 counts in at least one sample. These clusters were annotated with Ensembl GTF file version 87 annotations (downloaded May 2016), which includes known coding and noncoding RNA transcripts (**Supplemental Figure 2**). Unidirectional clusters overlapping known rRNA genes were also removed.

### CAGE Cluster Epigenetic and Transcription Factor Overlaps

Epigenetic datasets of interest were downloaded from their respective locations (**Supplemental Table 2**). Bam files were converted into tag directories using HOMER.^61^ HOMER annotatePeaks.pl was used to determine the read depth of each epigenetic dataset (normalized for cluster length and number) for each cluster annotation type (±1000bp of the cluster midpoint). HOMER findMotifsGenome.pl was used to check for enrichment of known transcription factors for each annotation type. For unidirectional clusters, the cluster midpoint ±200bp was used as input. For bidirectional clusters, the cluster boundaries were used and no additional nucleotides were added. Homer generated background sequences generated from a masked genome were used.

### Promoter Width Analysis

CAGEfightR was used to calculate the 0.1 to 0.9 inter-quantile range (IQR) for unidirectional clusters overlapping known promoter regions. A cutoff of 10bp was used to define a sharp and broad populations of promoters. The genomic coordinates of each promoter’s predominant TSS with 100bp added upstream and 50bp added downstream were inputted into bedtools getfasta to obtain genomic sequences.^62^ These sequences were inputted into the WebLogo tool to generate visual representations of nucleotide enrichment at each position relative to the predominant TSS.^63^ The genes of each promoter type were also inputted into the PantherGO online tool to check for enrichment of gene ontology terms ^64^. The R package ggplot2 was used to generate violin plots of sharp and broad promoter pooled expression levels and basepair width. To compare promoter IQR values across failed and healthy hearts, we first filtered out any promoters that were not present in all hearts. We used CAGEfightR with sample-specific scores to calculate the IQR of each promoter in each sample. We compared the average IQR across all promoters in the three healthy samples to the average IQR in the four failed samples using the nonparametric Wilcoxon rank sum test in R.

### Intronic Enhancer Analysis

A custom script was written to generate an annotation file of first intron locations for all transcripts present in the Ensembl GTF file version 87 (Downloaded May 2016). We used custom Python script to evaluate if bidirectional intronic completely overlapped the first intron of any transcript. ggPlot2 was used to generate violin plots of enhancer eRNA expression levels, base pair width, and bidirectionality scores. Genes with first intron and other intron enhancers were inputted into the PantherGO online software tool to check for enrichment of gene ontology terms.^64^

### RNA-Seq Data Analysis and Comparison with CAGE-Seq Data

Raw RNA-seq reads were trimmed with trimmomatic (v0.36) and aligned to the human genome (hg19) using STAR with default settings ^58^. Uniquely aligned reads were assigned to genes using htseq-count using the Ensembl GTF file version 87 (Downloaded May 2016) as annotations.^65^ Raw count matrices were inputted into EdgeR for normalization, dispersion estimation, and glm-model approach measures of differential expression between healthy and failed hearts.^66^ Genes with <1 count per million were removed from the analysis. We defined differentially expressed genes as any gene with an FDR-corrected p-value of < 0.05. The read counts of CAGE-seq unidirectional clusters overlapping gene promoters were used to quantify overall gene expression. Expression values from multiple promoters of the same gene were collapsed into a gene-level value. These count values were inputted into EdgeR and analyzed similar to the RNA-seq data above. ggPlot2 was used to graph the log-normalized and depth-normalized gene expression values generated by CAGE-seq and RNA-seq. The R package corrplot was used on the normalized count matrix to generate a correlation matrix across all samples.

Significantly downregulated and upregulated genes were separated based on the sign of their log fold-change value. Ensembl gene IDs were inputted into the PantherGO online tool to check for enrichment of gene ontology terms.^64^

### Differential Enhancer Analysis

Raw count values representing eRNA expression levels for bidirectional enhancers annotated as intragenic or intronic were exported as a counts table. This counts table was imported into EdgeR for differential expression analysis.^66^ Counts were normalized to library size, dispersion estimated, and differential enhancer usage called using glm-models. EdgeR was also used to generate an MDS plot of normalized enhancer counts. Due to the low count numbers associated with eRNA expression, expression values are subject to high levels of variation. EdgeR, like other RNA-seq analysis tools, detects this increased variation and reports higher p-values for calling differential expression. After multiple testing correction, there are too few enhancers surviving for downstream analysis. Therefore, raw p-value cutoffs were used. Enhancers with raw p-values ≤ 0.05 were used as input to HOMER findmotifsGenomewide.pl to find *de novo* motif enrichments in differential enhancers (enhancers with raw p-values > 0.05 were used as background sequences).^61^

### Alternative Promoter Usage Analysis

Unidirectional CAGE clusters overlapping annotated promoters were used as our promoter set. We required that an individual promoter make up at least 1% of total gene counts in all samples to be included in our analysis. A python script was written to count the number of promoters per gene. A python script was also written to calculate the percent usage of each promoter. The percent usage was averaged for the 3 healthy hearts and 4 failed hearts and the difference was calculated. To assess the alternative promoters’ effect on gene protein sequence, a custom annotation of all transcripts’ start codons was generated using the Ensembl GTF file version 87 (Downloaded May 2016).

### Enhancer Validation with Other Methods

Enhancer files from published sources were downloaded. To determine overlap with the Vista enhancer browser, enhancers with heart signal, enhancers with any positive signal, and enhancers with no signal were downloaded.^33^ For Dickel et al. 2016, the “Putative human heart enhancers identified by integrative analysis” table was used as enhancer predictions.^20^ For Spurrell et al. 2019, the “Predicted Enhancers in any 2 Samples” file was used as enhancer predictions. For FANTOM data, the “Ubiquitous enhancers organs” file for enhancer regions active in all organs tested was used. The FANTOM left ventricle and cerebellum enhancer sets were determined by requiring that the enhancer have non-zero expression in each tissue.^35^ As a negative control, genomic coordinates of enhancer regions were scrambled 500 times, avoiding placement in repeats or gap sequences. Negative control regions and CAGE sequence defined enhancer regions were intersected with downloaded predictions using bedtools intersect requiring at least 1bp overlap ^62^. Significance was determined using a fisher exact test.

## Acknowledgements

We thank the patients for their participation. We also thank Dr. Xinkun Wang from the Northwestern University sequencing core and Yujiro Takegami from DNAFORM for their excellent technical support. AG conducted the analysis and drafted the mansuscript. LDC secured patient consent and genotype information. PP assisted with genotyping. JAW provided access to control samples. DYB and MJP provided helpful advice and commentary and assisted with interpretation. MAN and EMM assisted with analysis, writing and editing the manuscript.

## SUPPLEMENTAL MATERIAL for Gacita et al

**Supplementary Table 1.** Sequencing Read Yields and Mapping Rates for CAGE-seq and RNA-seq Libraries.

| Sample | CAGE-Seq |  |  | RNA-Seq |  |
| --- | --- | --- | --- | --- | --- |
|  | pM | Total reads<br>(75bp SE) | Uniquely<br>Aligned Reads<br>(%) | Total Reads<br>(150bp PE) | Uniquely<br>Aligned<br>Reads (%) |
| Healthy<br>1 | 8.10 | 59,865,536 | 45,191,501<br>(75.49) | 53,274,910 | 41,113,982<br>(77.17) |
| Healthy<br>2 | 7.34 | 56,226,409 | 44,062,633<br>(78.37) | 58,609,578 | 44,742,094<br>(76.34) |
| Healthy<br>3 | 5.07 | 32,763,023 | 25,314,481<br>(77.27) | 49,620,253 | 38,276,453<br>(77.14) |
| Heart<br>Failure<br>1 | 5.30 | 39,295,214 | 30,906,318<br>(78.65) | 49,587,645 | 38,077,307<br>(76.79) |
| Heart<br>Failure<br>2 | 8.25 | 45,584,689 | 36,344,157<br>(79.09) | 43,147,599 | 32,813,912<br>(76.05) |
| Heart<br>Failure<br>3 | 8.03 | 46,398,662 | 35,734,514<br>(77.02) | 47,529,437 | 35,808,836<br>(75.34) |
| Heart<br>Failure<br>4 | 4.53 | 22,668,805 | 18,101,857<br>(79.85) | 40,737,687 | 31,579,044<br>(77.52) |

*pM*, picomolar. *SE*, single end. *PE*, paired end

**Supplementary Table 2.** Epigenetic Datasets used for CAGE-Cluster Functional Annotation

| **Target** | **Assay Type** | **Tissue Source** | **Accession #** |
| --- | --- | --- | --- |
| Open Chromatin | ATAC-Seq | Female adult (51 years) LV Tissue | ENCSR117PYB |
| Open Chromatin | DNase-Seq | Female adult (53 years) LV Tissue | ENCFF702IJE |
| H3K4me1 | ChIP-Seq | Male child (3 years) LV Tissue | ENCFF901JPP |
| H3K4me3 | ChIP-Seq | Male Adult (34 years) LV Tissue | ENCFF527LGE |
| H3K27Ac | ChIP-Seq | Female Adult (51 years) LV Tissue | ENCFF625XET |
| CTCF | ChIP-Seq | Female Adult (53 years) LV tissue | ENCFF738KRH |
| POL2A | ChIP-Seq | Female Adult (53 years) LV tissue | ENCFF318MWF |

*ATAC*, assay for transposase-accessible chromatin. *ChIP*, chromatin immunoprecipitation.

### Supplementary Figures

**Supplementary Figure 1.**
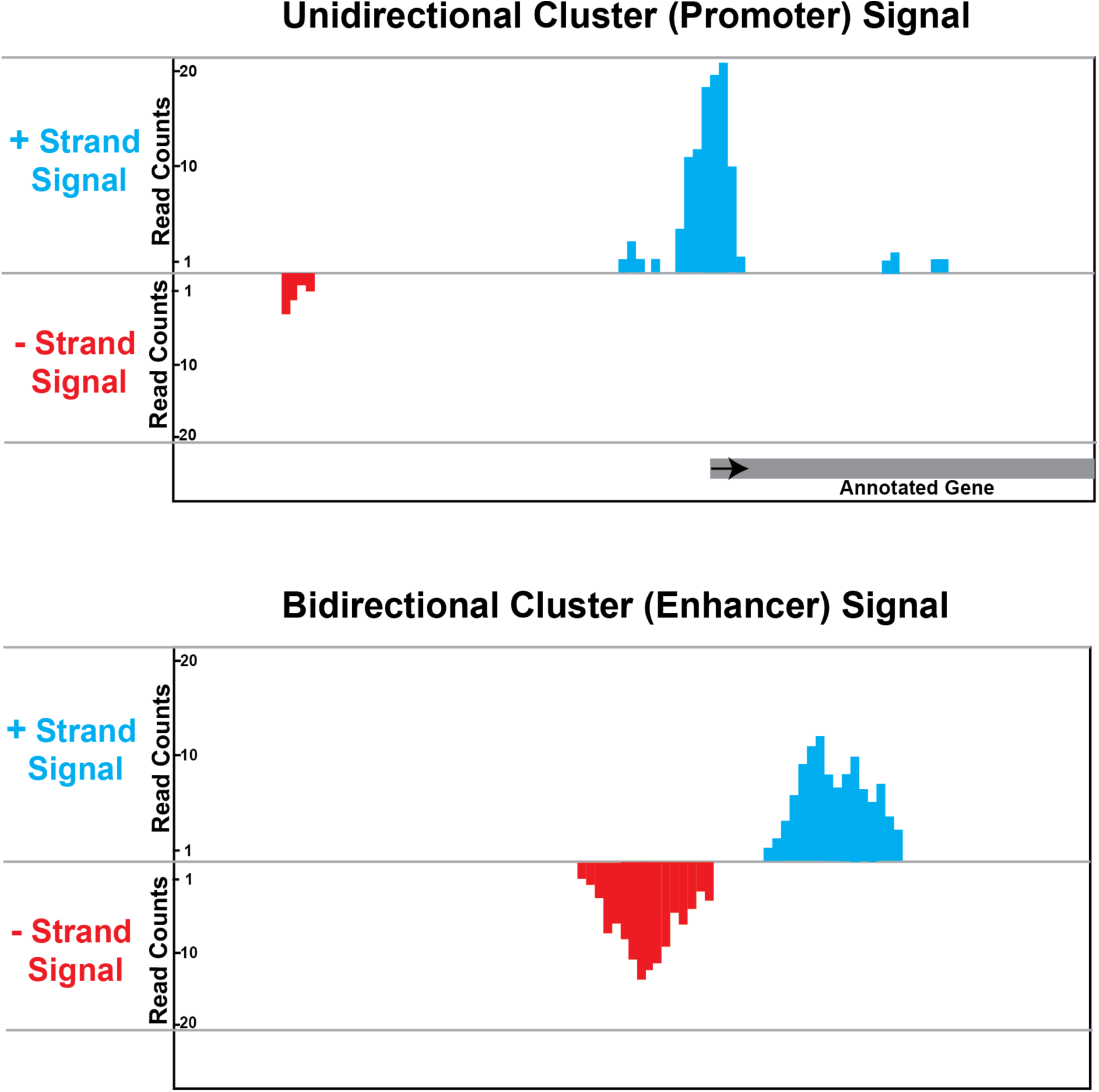
Example schematic of unidirectional (top) and bidirectional (bottom) CAGE clusters representing promoters and enhancer regions, respectively. Positive (sense) strand signals are shown in blue and minus (antisense) signals are shown in red.

**Supplementary Figure 2.**
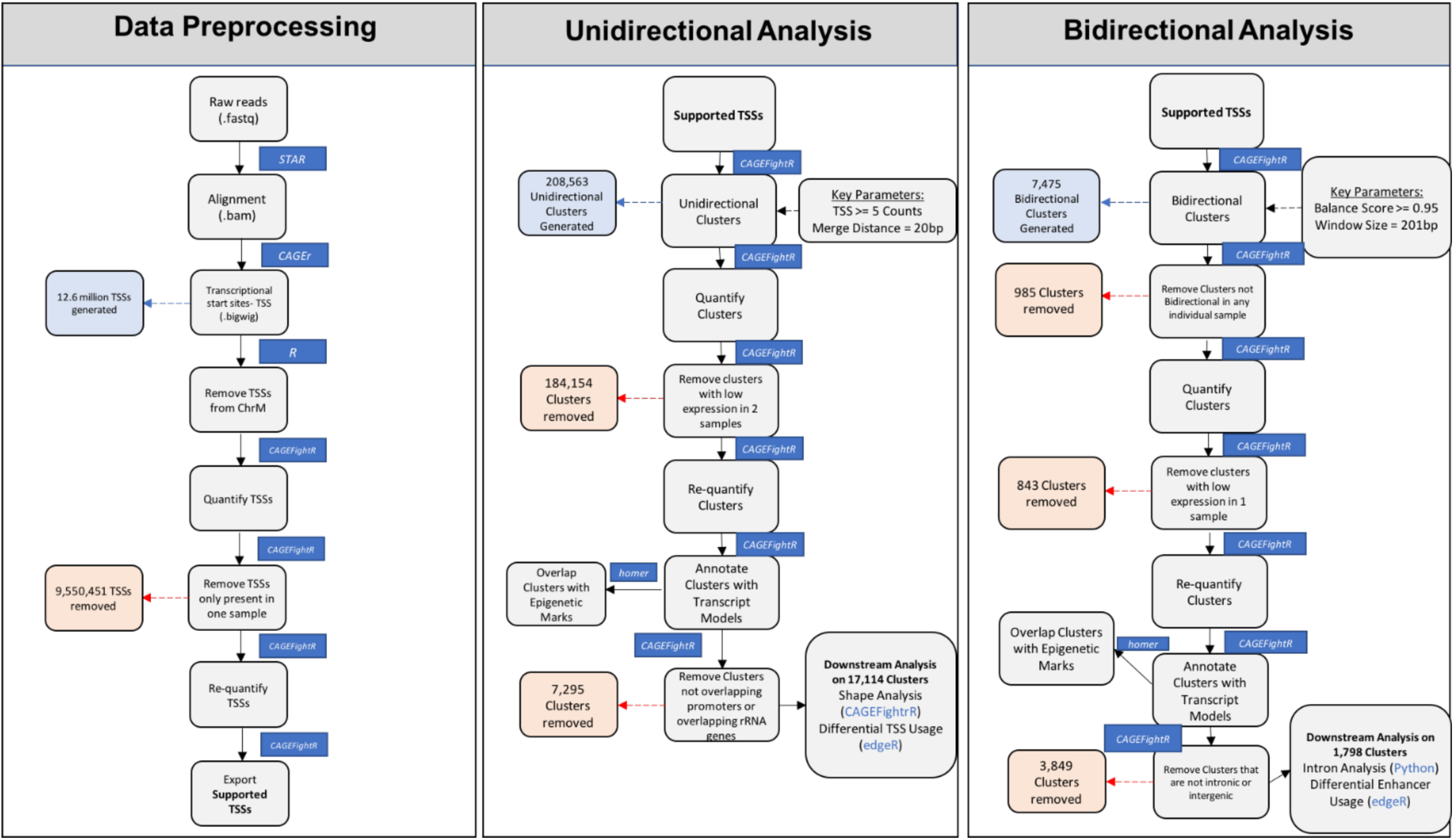
Data analysis pipeline for identifying and analyzing promoters and enhancers from CAGE-seq data. *TSS*, transcriptional start sites.

**Supplementary Figure 3.**
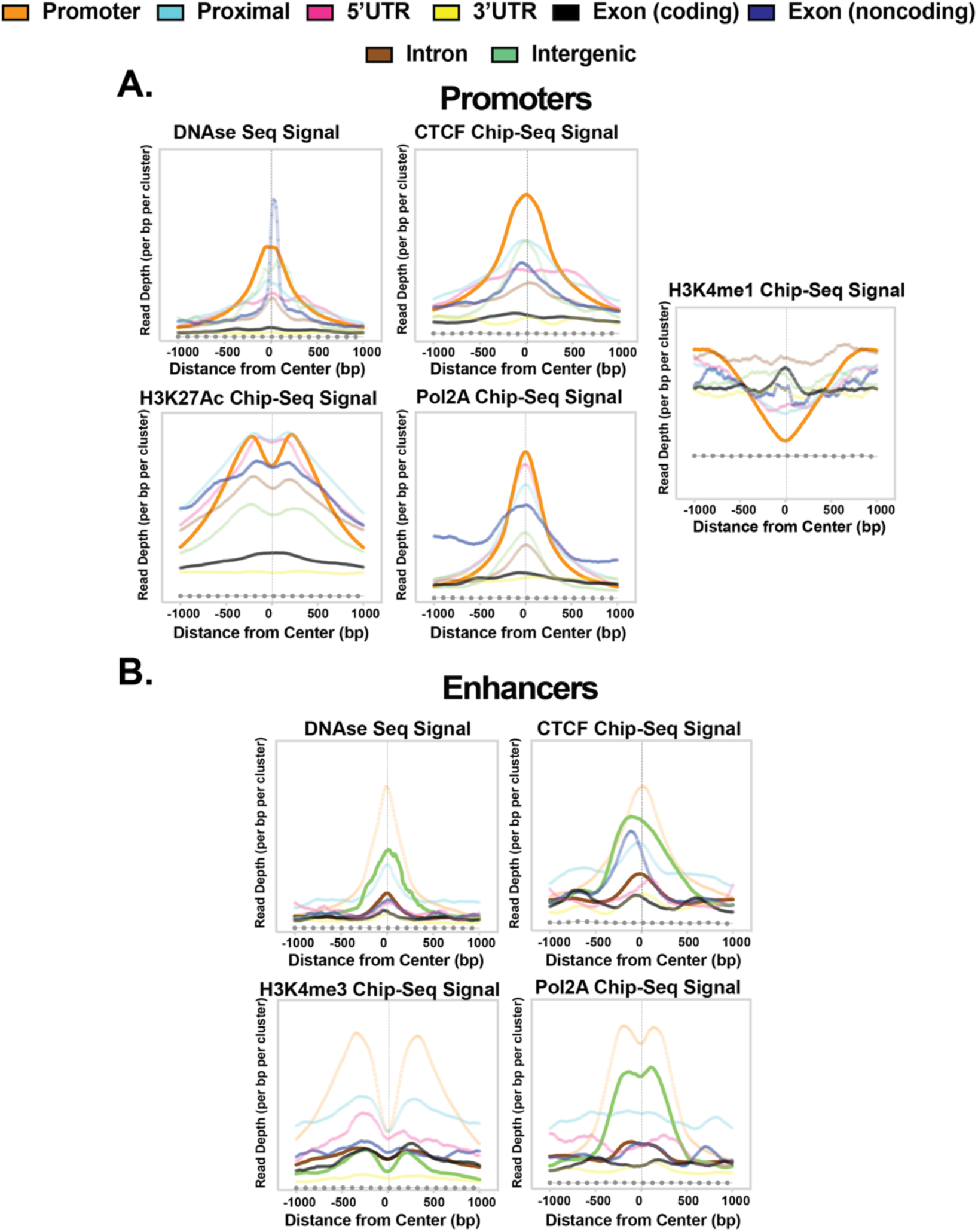
Additional left ventricle epigenetic signals of unidirectional and bidirectional CAGE clusters. **A.** Left ventricle open chromatin (DNAse-seq), protein binding (CTCF, Pol2A), and enhancer histone marks (H3K27Ac and H3K4me1) signals for unidirectional CAGE clusters of all annotation classes. **B.** Left ventricle open chromatin (DNAse-seq), protein binding (CTCF, Pol2A), and promoter histone marks (H3K4me3) signals for bidirectional CAGE clusters. Dashed lines in A and B represent signals from genomic regions created by scrambling the location of unidirectional and bidirectional clusters, respectively.

**Supplementary Figure 4.**
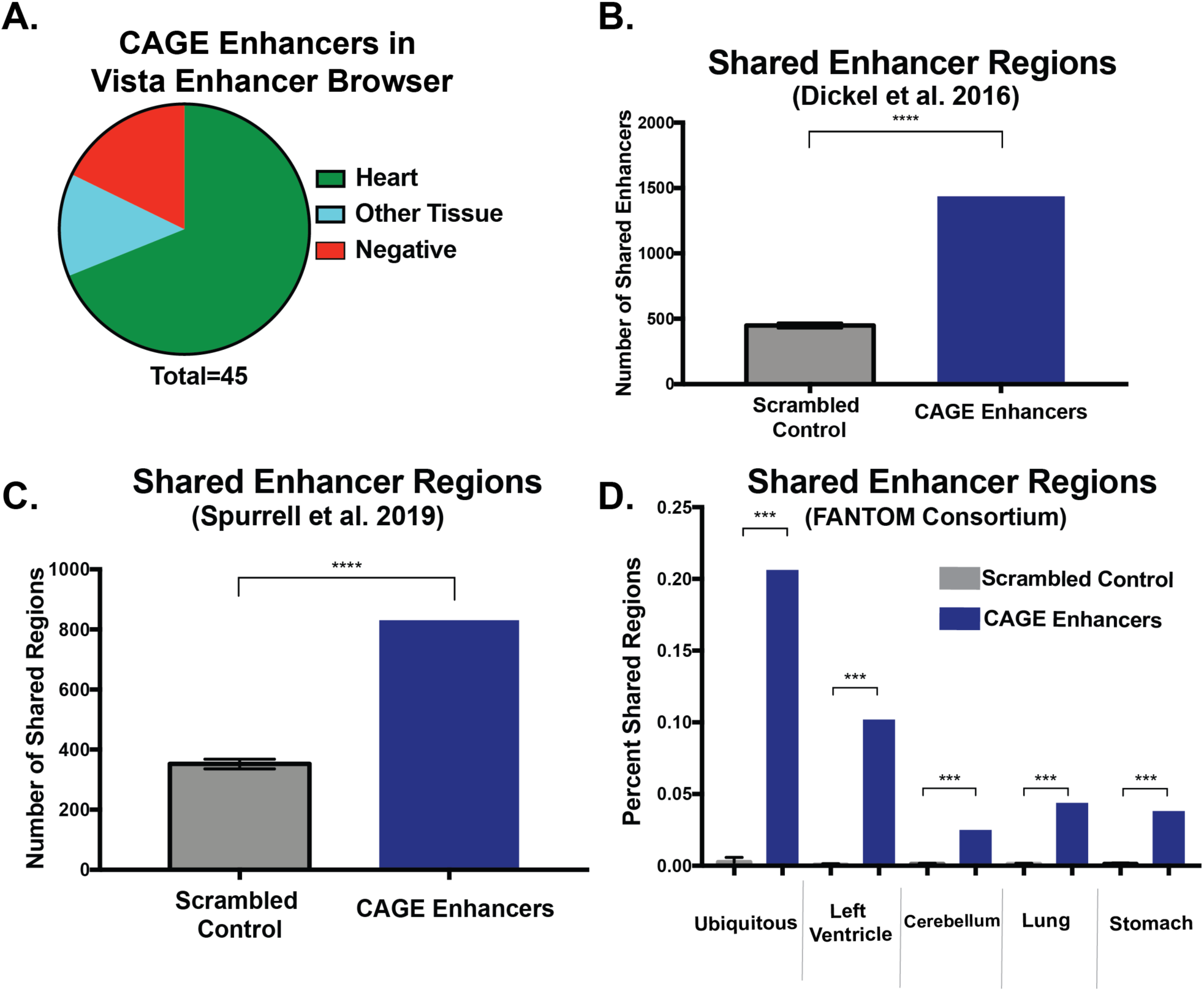
CAGE-defined enhancers overlapped enhancers determined by independent methods. **A.** Venn diagram of Vista Enhancer Browser data displaying the results of functional testing of 45 CAGE enhancers. **B & C**. Bar charts showing the number of overlapping enhancers when comparing CAGE and two independent methods. **D**. Bar charts indicating the percentage of FANTOM enhancers that overlap CAGE enhancers for five different groups of FANTOM enhancers. Scrambled controls represent the number of overlaps obtained when randomly shuffling the genomic location of CAGE enhancers. Significance determined by fisher’s exact test. (p ≤ 0.05(*), ≤ 0.005(**), ≤ 0.0005 (***)).

